# Cocaine and caffeine elicit different dopamine receptor-mediated locomotor activity profiles: insights from a new model

**DOI:** 10.64898/2026.01.12.699083

**Authors:** Jordan Moass, Krithik Ashokkumar, Martin O. Job

**Author notes:** Corresponding author: (MOJ).

## Abstract

**Background:** The locomotor activity elicited by cocaine and caffeine can be distinguished via their underlying mechanisms and via differential responses to neuropharmacological manipulations. However, both cocaine- and caffeine-induced locomotor activity can be blocked by non-selective dopamine receptor antagonists, implying that we may not be able to distinguish their locomotor activity at the level of the dopamine receptor-mediated mechanism. With the rationale that this limitation may be due to a lack of sensitivity of current methods, we have developed a new Quantitative Structure of Curve Analytical (QSCAn) model. We hypothesized that QSCAn will be more effective, relative to the current model, in differentiating the dopamine receptor-mediated mechanism of cocaine versus caffeine-induced locomotor activity.

**Methods:** We assessed locomotor activity (quantified as distance traveled in cm over time) due to injections of saline (n = 6), cocaine (10 mg/kg i.p, n = 8) and caffeine (20 mg/kg i.p, n = 6) in male Sprague Dawley rats with and without pretreatment with vehicle and cis-flupenthixol (0.2 mg/kg i.p, non-selective dopamine receptor blocker)(current model). Because distance traveled over time follows an inverted u-shaped time-response curve, we employed gaussian fit of this structure to obtain several behavioral variables (QSCAn model). We compared both models.

**Results:** The current model could not distinguish the locomotor activity profile of cocaine versus caffeine following vehicle and cis-flupenthixol pretreatments. QSCAn model could distinguish cocaine versus caffeine in the presence and absence of non-selective dopamine receptor blockade.

**Conclusions:** The new QSCAn model may be a promising tool to distinguish/characterize different psychostimulant-related effects.

## Introduction

Psychostimulants include illicit substances such as cocaine and amphetamines, and legal substances such as caffeine and tea. In rodent models, administration of these psychostimulants result in increases in locomotor activity (Antoniou et al., 1998). This increase in locomotor activity is thought to be related to increases, in part, in dopaminergic signals in the brain, though the underlying mechanisms remain incompletely understood. Cocaine inhibits the dopamine transporter to increase dopamine which in turn acts at the dopamine receptors to increase locomotor activity (Fig 1). Caffeine is an adenosine receptor antagonist which disinhibits the dopamine receptor component of an adenosine-dopamine receptor heteromer to amplify dopamine effects which in turn can increase locomotor activity (Fig 1). There is evidence that not only do caffeine and cocaine have different mechanisms of action (Fig 1), but the locomotor activity elicited by cocaine and caffeine can be distinguished via their differential responses to neuropharmacological manipulations (Table 1).

**Fig 1:**
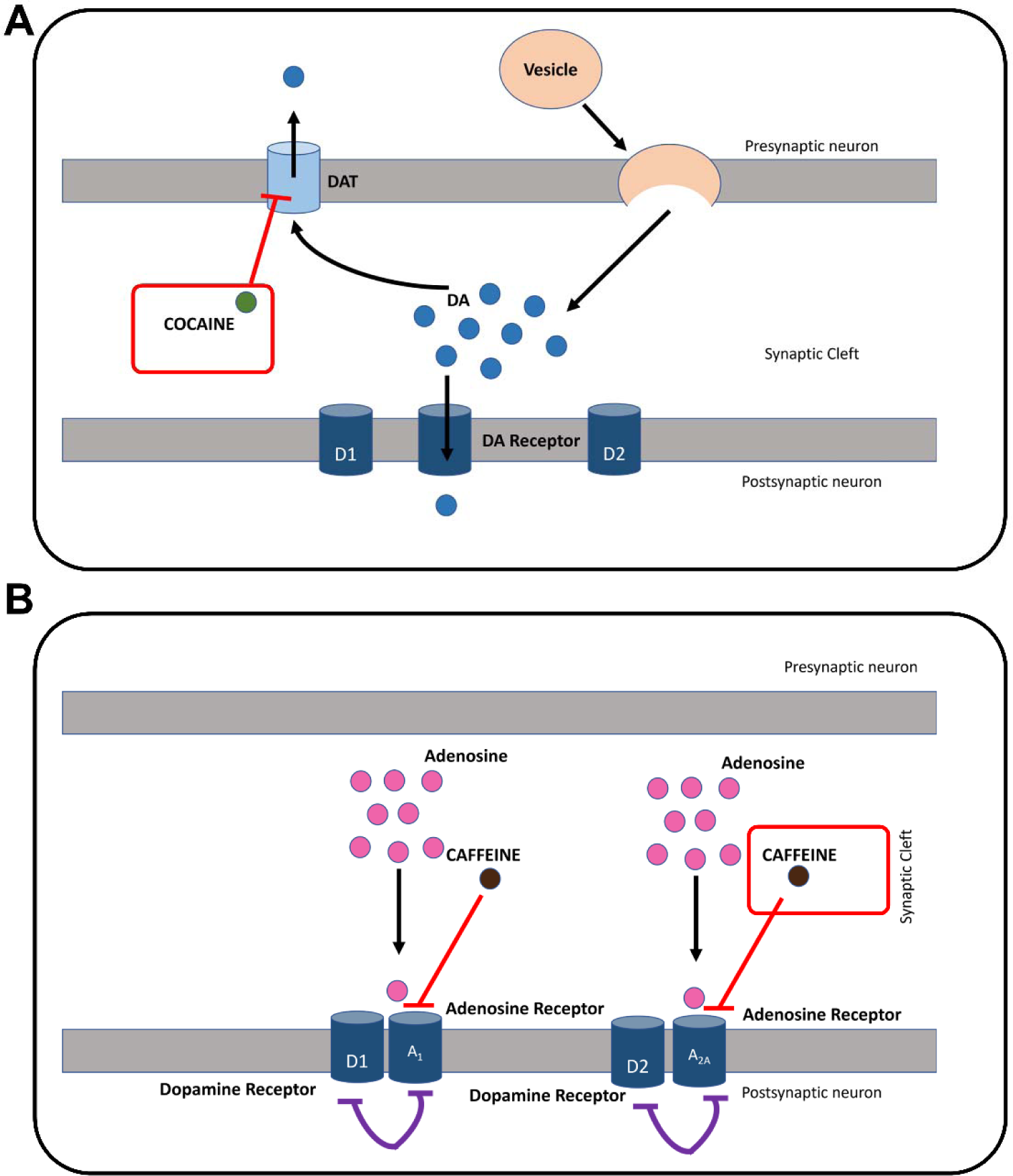
Differences in the proposed mechanism of action of cocaine versus caffeine on dopamine-mediated locomotor activity. Cocaine inhibits the dopamine transporter causing an increase in extracellular dopamine which acts at dopamine D1 and D2 receptors resulting in increases in locomotor activity. Caffeine inhibits the adenosine A1 and A2 receptors to relieve adenosine-mediated inhibition of dopamine D1 and D2 receptors, respectively, increasing locomotor activity with or without increases in extracellular dopamine levels.

**Table 1:** Cocaine- and caffeine-induced locomotor activity profiles appear to be different, but it is unclear if these psychostimulants elicit different behavioral outputs at the level of the dopamine receptor.

| Route of administration, drug | cocaine | caffeine |
| --- | --- | --- |
| Intra-nucleus accumbens, CART peptide | Decrease (Job, 2016) | No effect (Job, 2016) |
| Intra-nucleus accumbens, NMDA antagonist | Decrease (Pulvirenti et al., 1991) | No effect (Pulvirenti et al., 1991) |
| Intra-nucleus accumbens, glutamate antagonist | Decrease (Pulvirenti et al., 1989) | No effect (Pulvirenti et al., 1989) |
| Systemic NMDA antagonist | Decrease (Karler and Calder, 1992) | No effect (Powell and Holtzman, 1998) |
| Intracerebroventricular, intra-nucleus accumbens, neurotensin | Decrease (Robledo et al., 1993) | No effect (Skoog et al., 1986) |
| Intracerebroventricular, CRF antibody | Decrease (Sarnyai et al., 1992) | No effect (Sarnyai et al., 1992) |
| Systemic, dopamine antagonist | **Decrease** (Baldo et al., 1999; Delfs et al., 1990; Kovács et al., 1990; Wenzel et al., 2013) | **Decrease** (Garrett and Griffiths, 1997; Garrett and Holtzman, 1994; Koob et al., 1984) |

Interestingly, while cocaine- and caffeine-induced locomotor activity can be distinguished in their responsivity to pharmacological of several neurochemical/ neurohormonal systems, it is unclear that they can be distinguished via their responses to pharmacological blockade of the dopamine receptor (Table 1). The effects on the locomotor activity of these psychostimulants using dopamine lesions/ablation are equivocal: no effect (French, 1986; French and Vantini, 1984; Joyce and Koob, 1981; Swerdlow and Koob, 1985) and decreases (Erinoff et al., 1984; Erinoff and Snodgrass, 1986; Estler, 1979; Finn et al., 1990) in caffeine-induced locomotor activity have been observed using these methods. But blockade of dopamine receptors suppresses caffeine-induced locomotor activity (Garrett and Griffiths, 1997; Garrett and Holtzman, 1994; Koob et al., 1984), not unlike cocaine (Table 1).

Based on the significant differences between these two psychostimulants mechanistically (Fig 1) and behaviorally (Table 1), we rationalized that an explanation for the lack of distinction in how cocaine and caffeine respond to dopamine antagonists may be related to the sensitivity, or the lack thereof, of current locomotor activity assessment models. With our goal being to advance the field, we reviewed the current methods of locomotor activity assessment. While doing this, we observed that distance traveled (a measure of locomotor activity) increases and then decreases back to baseline over time (Fig 2A). Because this ‘increase and decrease’ is reminiscent of an inverted u-shaped time-response (IUTR) curve structure (Fig 2B), we developed a new model, which we call the Quantitative Structure of Curve Analytical (QSCAn), to analyze this IUTR curve structure.

**Fig 2:**
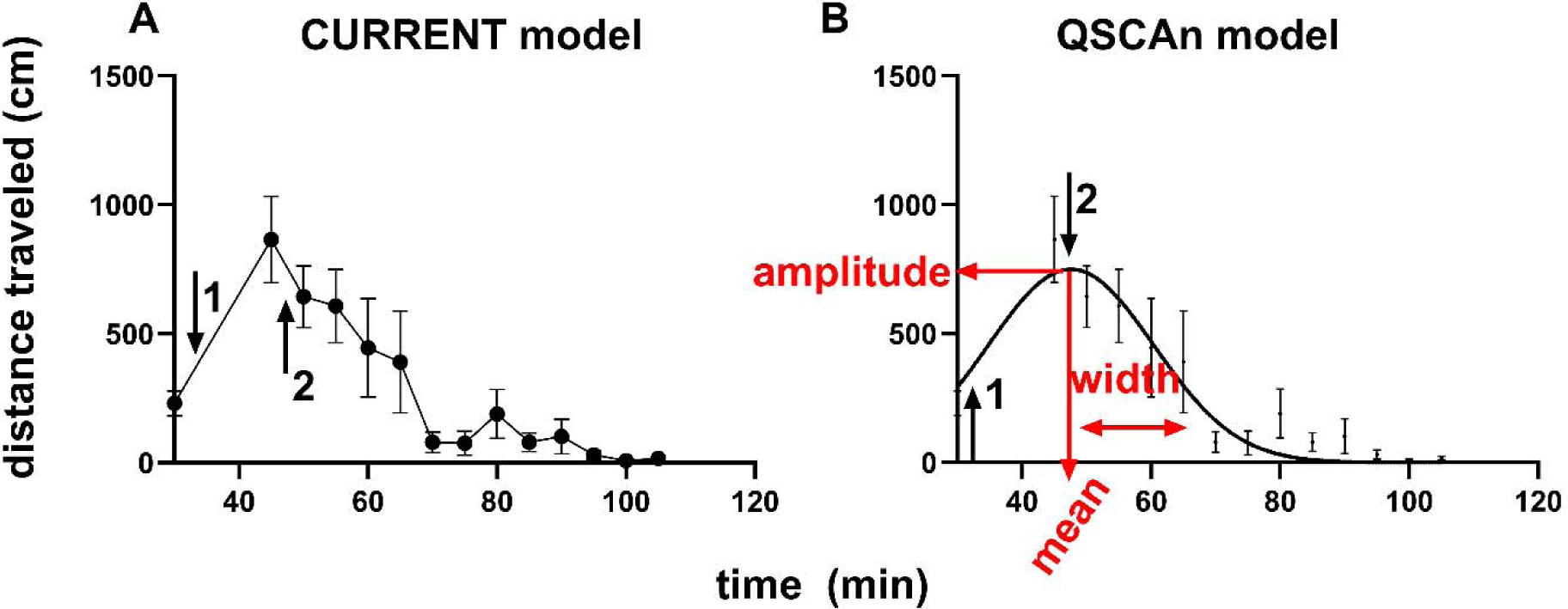
The current model and the Quantitative Structure of Curve Analytical (QSCAn) model. These graphs represent locomotor activity due to saline injection. Both models (current model in A and QSCAn model in B) are represented as plots of the same data with time on the x-axis and distance traveled (cm) on the y-axis. For both graphs, the arrows 1 and 2 represent, respectively, the injection time points for pretreatment (vehicle, cis-flupenthixol) and treatment (saline). Analysis of the current model graph includes repeated measures ANOVA with the dependent variable being distance traveled. We developed a new model for analysis of the time course data – shown in B. The idea behind the QSCAn model is based on the structure of the graph in A: distance traveled increases and then decreases as time elapses giving rise to an inverted u-shaped time-response curve (IUTR). For the QSCAn model analysis, we employed gaussian function fit for the IUTR curve structure of locomotor activity to reveal several variables including amplitude, mean, width. The amplitude is the maximum response. The mean is the time value at which the amplitude is maximum. The width is the distance from the mean to the time when the amplitude = 0. The goal of this study was to compare the current model and the QSCAn model for effectiveness in distinguishing the locomotor activity of cocaine versus caffeine. We hypothesize that QSCAn will reveal behavioral structure variables that can enable us to differentiate between cocaine and caffeine-related locomotor activity.

To quantify the variables that define an IUTR (see variable definitions in Fig 2), we can employ the gaussian fit (equation 1)

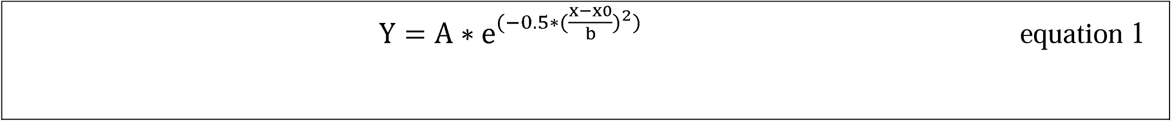

Where Y = distance traveled, X = time, A = amplitude, x0 = mean, b = width.

We hypothesized that our new QSCAn model (for assessment of the IUTR) will be more effective, relative to the current model, in differentiating cocaine-versus caffeine-induced dopamine receptor-mediated locomotor activity.

Both cocaine and caffeine-induced locomotor activity have a dopamine receptor-mediated component (Fig 1). To identify the dopamine receptor-mediated component of both cocaine versus caffeine-induced locomotor activity, we utilized the non-selective dopamine receptor antagonist – cis-flupenthixol which has been shown to block the locomotor activity of both of these psychostimulants (Baldo et al., 1999; Koob et al., 1984; Swerdlow and Koob, 1985; Wenzel et al., 2013). We designed experiments to test our hypothesis by comparing the current model versus QSCAn model (Fig 2A versus 2B) for cocaine-versus caffeine-induced dopamine-mediated locomotor activity. We wanted to know if we could detect differences between cocaine and caffeine locomotor activity profiles in the presence and absence of dopamine receptor blockade.

For the experiments, we assessed, in male Sprague Dawley rats, locomotor activity at 1) baseline, 2) after vehicle or cis-flupenthixol pretreatment, just before 3) saline, cocaine and caffeine treatment. For an effective dose that could block both cocaine and caffeine, we used cis-flupenthixol (0.2 mg/kg i.p, in counterbalanced design: all subjects received both pretreatments on different experimental days) prior to injections of saline, cocaine and caffeine and we assessed locomotor activity (as distance traveled over time). We compared the current model and the QSCAn model to determine which model could differentiate cocaine versus caffeine locomotor activity profiles while blocking or not blocking dopamine receptors. Our methods, results, discussion and conclusion are below.

## Methods and Materials

### Subjects

Experiments and animal care were in accordance with the Institute of Animal Care and Use Committee of Emory University and the National Institutes of Health Guide for the Care and Use of Laboratory Animals. A total of twenty (20) male Sprague Dawley rats (Charles River Inc., Wilmington, MA) were used. They were provided with rat chow and water ad libitum and maintained on a 12 h light: dark cycle (lights on at 7am). We conducted all experiments between 10:00 AM and 6:00 PM. For the experiments, the animals weighed 400 – 700 g.

### Animal Health

Throughout the experiments from the arrival of the rats at the housing facility through the end of behavioral experiments, we ensured that rats were not in any pain or distress. After the completion of all behavioral experiments, the rats were euthanized via isoflurane anesthesia followed by decapitation according to established guidelines.

### Locomotor activity measurement equipment

We assessed locomotor activity in locomotor chambers (Omnitech Electronics, Columbus OH). These locomotor chambers included transparent Plexiglas walls (dimensions of 40 × 40 × 30 cm) and a photocell cage containing 32 photobeams located 5 cm above the floor (Omnitech Electronics, Columbus OH). The experimental apparatus was such that each photocell cage was connected to a computer equipped with software (Digipro; Omnitech Electronics) to measure locomotor activity.

### Habituation to test procedures

For three days before the experiment, we placed the rats into locomotor chambers for 30 min to allow them to be habituated to their test environment. At this time also, we habituated rats to injection procedure – we injected with saline via the intraperitoneal (i.p) route after removing them from the chambers. After injection, we returned them into the chambers, but no data was obtained. On the experiment day, we placed the animals for 30 min into the locomotor chambers. After the 30 min, we started obtaining locomotor activity data.

### Experimental procedure

Locomotor activity assessments are as previously described (Job, 2016; Job et al., 2014, 2013, 2012; Job and Kuhar, 2012). For the systemic studies, rats were placed into locomotor chambers for 30 min to allow them to be habituated to their surroundings before the recording of basal locomotor activity commenced. Locomotor activity was assessed as distance traveled in centimeters (cm). After 30 min of basal locomotor activity recording, rats were removed from the chambers and pretreatments (vehicle and cis-flupenthixol 0.2 mg/kg i.p) were administered. After pretreatments, subjects were placed back into the chambers and locomotor activity was recorded for 15 minutes. After we obtained the locomotor activity data for 15 min, the rats were (again) removed from the locomotor chambers and administered psychostimulants (saline as control, cocaine or caffeine) via the intraperitoneal route and placed back into the chambers for locomotor activity measurements for an additional 60 min. On the different experiment days (vehicle and cis-flupenthixol pretreatment experiment sessions), rats were placed in the same locomotor chamber.

### Psychostimulants (drugs and drug doses)

Cocaine hydrochloride was from NIDA. Caffeine [1,3,7-Trimethyl xanthine] was from Sigma-Aldrich, St Louis, MO. α-flupenthixol dihydrochloride [cis-Z-Flupenthixol dihydrochloride] was obtained from Sigma-Aldrich, St Louis, MO. All systemic injections were given in volumes of 1 mL/ kg. Injection doses for cocaine and caffeine were 10 mg/kg i.p and 20 mg/kg i.p, respectively. Injection doses for cis-flupenthixol were 0.0 (vehicle) and 0.2 mg/kg via the i.p route.

### Rationale for drug doses

We chose 10 mg/kg of cocaine because we have shown in several studies that this dose increases locomotor activity significantly above baseline locomotor activity (Job et al., 2014, 2013, 2012; Job and Kuhar, 2017; Kuhar and Job, 2017). For caffeine, we utilized the 20 mg/kg dose because this dose increases locomotor activity above baseline locomotor activity levels (Job, 2016).

The dose of cis-flupenthixol that we used was 0.2 mg/kg. This dose attenuates locomotor activity due to amphetamine (Koob et al., 1984; Powell and Holtzman, 1998; Swerdlow and Koob, 1985; Vaccarino et al., 1986), cocaine (Baldo et al., 1999), and caffeine-induced locomotor activity (Koob et al., 1984; Swerdlow and Koob, 1985). Furthermore, it blunts the locomotor activity driven by opioid injections into ventral tegmental area (Joyce et al., 1981)-we interpret this to imply that this dose is effective in suppressing dopamine-mediated locomotor activity.

Specifically, 0.25 mg/kg dose of cis-flupenthixol has been shown to effectively blunt cocaine (15 mg/kg i.p)-induced locomotor activity (Wenzel et al., 2013) but not locomotor activity driven by glutamatergic activity (Tuplin et al., 2015) - we interpret this to imply that this dose is effective for dopamine-mediated but not glutamate-mediated locomotor activity. At a cis-flupenthixol dose of 0.3 mg/kg locomotor activity and also feeding behavior are suppressed (Pitts and Horvitz, 2000) – we interpret this to imply that this dose is too high. We chose the 0.2 mg/kg dose of cis-flupenthixol as a dose that blocks dopamine-mediated locomotor activity irrespective of the drug administered (caffeine or cocaine).

### Experimental Design

We conducted experiments to determine the effects of systemic administration of cis-flupenthixol on locomotor activity due to systemic administration of saline, cocaine and caffeine. Experiments were conducted in counterbalanced design to ensure that each subject was its own control, with at least three days between experiments. Different subjects were used for different psychostimulant treatments (saline, cocaine and caffeine), but within each psychostimulant treatment the same subjects were administered both of the pretreatments (cis-flupenthixol 0.0 mg/kg i.p (saline) and cis-flupenthixol 0.2 mg/kg i.p). See the experiment designs in Fig 3.

**Fig 3:**
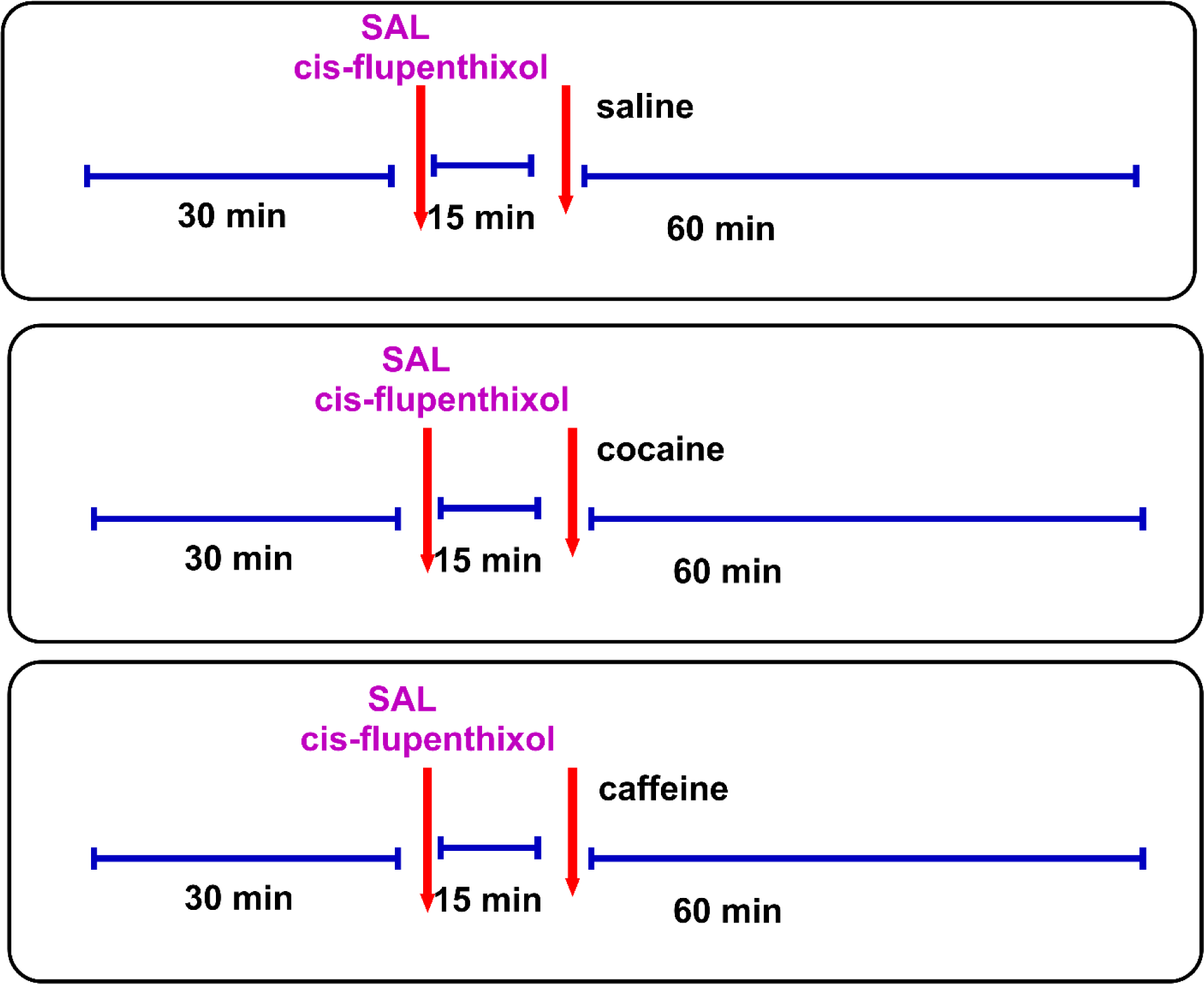
Experimental design: After 30 min of baseline locomotor activity assessment, male rats (n = 20) were injected with vehicle (saline) and cis-flupenthixol (0.2 mg/kg i.p) and 15 min later, they were injected with saline, cocaine (10 mg/kg i.p) or caffeine (20 mg/kg i.p) followed by locomotor activity assessments for 60 min. Each subject received both vehicle and cis-flupenthixol in counterbalanced design on different experiment days. Different groups of rats were injected with saline (n = 6), cocaine (n = 8) or caffeine (n = 6).

### Locomotor activity data obtained

As shown in Fig 3, the experiment design has 3 components – 1) baseline (30 min, 5 min bins, 6 sample collected), 2) pretreatment (15 min, 5 min bins, 3 samples), and 3) treatment (60 min, 5 min bins, 12 samples). For analysis, for each subject, we averaged the distance traveled in the first 30 min and made this a single time point at time = 30 min. We also averaged the distance traveled in the next 15 min and made this a single time point at time = 45 min. All other distances were used as obtained (per subject). For data analysis, we used 14 time points per subject.

### Power Analysis

We conducted *A priori* power analysis using G*Power 3.1.9.7 for ANOVA-repeated measures within-between interaction using the following values: effect size = 0.6, α error prob = 0.05, power (1-ß err prob) = 0.95, number of groups = 3, number of measurements = 14, correlation among repeated measures = 0.5, Nonsphericity correlation = 1. This yielded a sample size N = 6 per group.

### Statistical Analysis

Analyses were performed using GraphPad Prism version 10 (GraphPad Software Inc, La Jolla, CA), SigmaPlot for Windows version 14.5 (Systat Software Inc, San Jose, CA), and JMP Pro version 18 (SAS Institute Inc., Cary, NC. Data were all expressed as mean ± SEM, and when significance occurs is set at P < 0.05. Normal distribution of all data was confirmed using Shapiro-Wilk tests (W, data set is determined to pass normality test when P > 0.05). Constant variant test was conducted using Spearman’s rank correlation and was determined to fail if P < 0.05. We employed Grubb’s test in GraphPad with P < 0.05 to determine if there were outliers within the data set.

For the current model, we compared the locomotor activity time course of saline, cocaine and caffeine with and without dopamine receptor blockade. For each pretreatment separately (vehicle, cis-flupenthixol), we employed Two-way repeated measures ANOVA with factors being drug (saline, cocaine, caffeine) and time (14 levels).

For the QSCAn model, we employed gaussian fit (see equation 1) of the *same* time course graphs as the current model (graph of distance traveled (cm) on y-axis and time on the x-axis) to derive an IUTR curve structure that is defined by several variables (amplitude, mean, width, AUC, see Fig 2). Using non-linear regression analysis, we compared the IUTR curve structures of saline, cocaine and caffeine with and without pretreatment of the dopamine receptor blocker.

For the current model, we compared the effects of dopamine receptor blockade versus vehicle control on the locomotor activity time course of saline, cocaine and caffeine (separately). For each drug (saline, cocaine and caffeine) separately, we employed Two-way repeated measures ANOVA with factors being pretreatment (vehicle, cis-flupenthixol) and time (14 levels). For the QSCAn model, we compared the IUTR curve structures of the locomotor activity time course following vehicle versus cis-flupenthixol for saline, cocaine and caffeine.

## Results

### The Methods we employed are described above. Grubb’s test detected no outliers

For both models (see Fig 2), we plotted distance traveled on the y-axis and time on the x-axis. We analyzed the current model (Fig 2A) using Two-way repeated measures ANOVA while we analyzed the QSCAn model using non-linear regression analysis (with gaussian fit) (Fig 2B).

### Vehicle pretreatment: saline versus cocaine versus caffeine

For experiments that included vehicle pretreatment (Fig 4A-B), Two-way repeated measures ANOVA with factors drug (saline, cocaine, caffeine) and time (14 levels) revealed the following: drug × time interaction (F 5.846, 49.69 = 3.943, P = 0.0029), a main effect of drug (F 2, 17 = 5.973, P = 0.0108), and a main effect of time (F 2.923, 49.69 = 7.884, P = 0.0002). Tukey’s post hoc test revealed that the time course for saline and cocaine/caffeine were different from time = 65 min to the end (* for cocaine and $ for caffeine in Fig 4A). However, Tukey’s post hoc tests revealed no significant differences between cocaine and caffeine at all/any time points (Fig 4A). Using this same current analytical model, we wanted to compare (without saline) the psychostimulants (cocaine and caffeine). Two-way repeated measures ANOVA revealed the following results: no psychostimulant × time interaction (F 2.609, 31.31 = 1.881, P = 0.1593), no main effect of psychostimulant (F 1, 12 = 1.258, P = 0.2841), but a main effect of time (F 2.609, 31.31 = 8.358, P = 0.0005). In summary, under vehicle pretreatment, the current model did not detect any differences between the distance traveled-time course of caffeine and cocaine.

**Fig 4:**
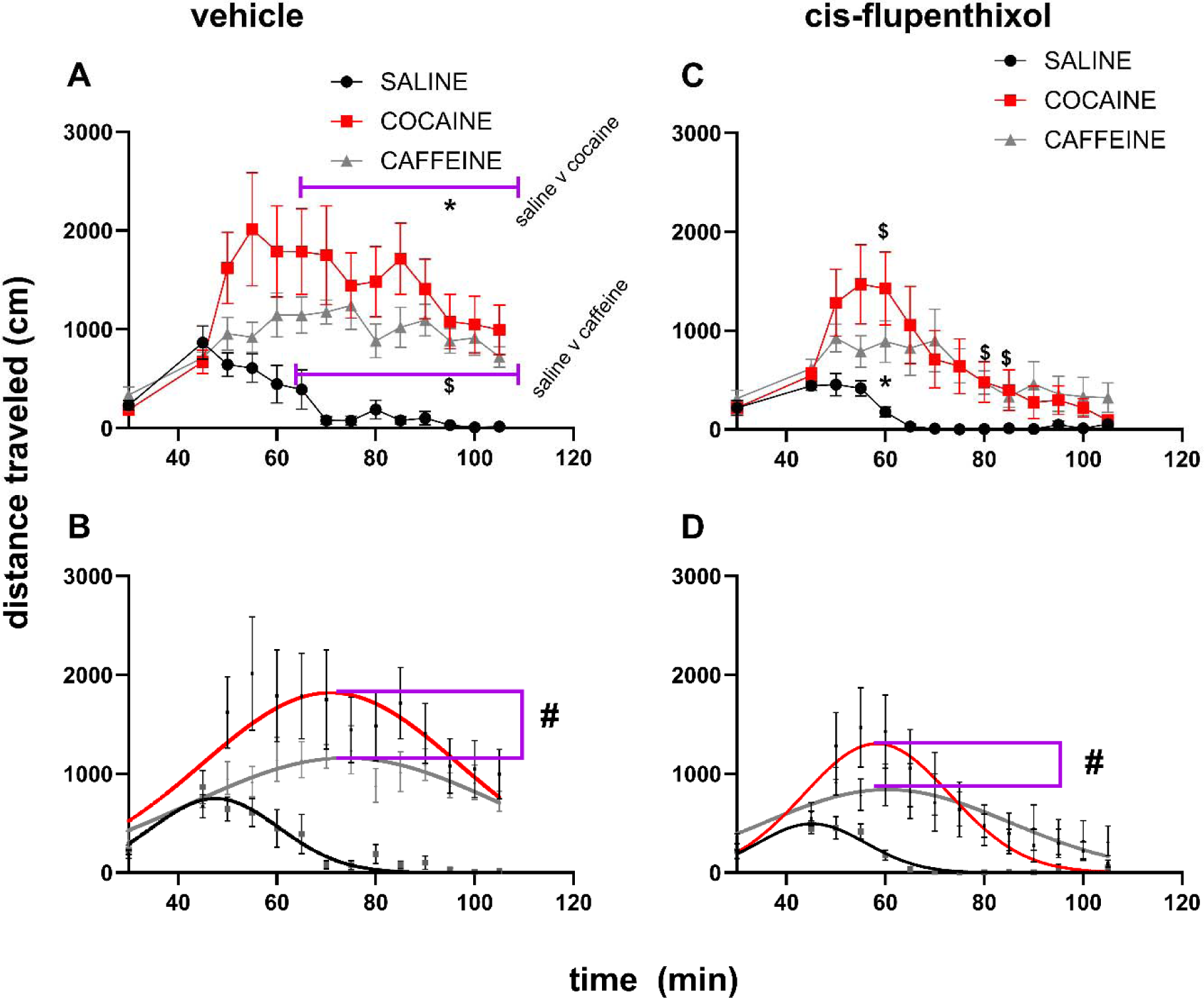
The Quantitative Structure of Curve Analytical (QSCAn) model was a more effective tool for the purposes of distinguishing the response-time course of cocaine versus caffeine-related locomotor activity following vehicle and cis-flupenthixol pretreatment. Black, red, and gray symbols and lines represent saline, cocaine, and caffeine treatment groups, respectively. A represents the time-response graph for the groups treated with saline prior to treatments with saline, cocaine and caffeine. B represents A when expressed as the QSCAn model. C and D represent, respectively, the current and QSCAn model time-response graphs for the groups treated with cis-flupenthixol prior to treatments with saline, cocaine and caffeine. Under vehicle pretreatment, Two-way repeated measures ANOVA showed no differences between cocaine and caffeine at any time point (current model, A) whereas a comparison between the structures defined by amplitude, mean and width (all together) for saline, cocaine and caffeine IUTR showed significant differences (P < 0.0001, B). Under cis-flupenthixol pretreatment, Two-way repeated measures ANOVA showed no differences between cocaine and caffeine at any time point (current model, C) while a comparison between the structures defined by amplitude, mean and width (all together) for saline, cocaine and caffeine IUTR showed significant differences (P < 0.0001, D). Under both vehicle and cis-flupenthixol pretreatment conditions, we detected differences between the IUTR of cocaine versus caffeine (C, D). In summary, in the absence (A-B) and presence (C-D) of non-selective dopamine receptor blockade, the QSCAn model detected differences between caffeine and cocaine-related locomotor activity whereas the current model did not.

For this same time course data (Fig 4A), we fitted the distance traveled-time course data using gaussian fit to derive an IUTR curve structure (Fig 4B, QSCAn model). A table of the variables of the IUTR structure and the adjusted R^2^ values are shown in Table 2. A comparison between saline, cocaine and caffeine IUTR showed significant differences (F 6, 271 = 24.92, P < 0.0001, Fig 4B). For comparisons between saline, cocaine and caffeine locomotor activity structure with respect to the individual variables, we obtained the following results: amplitude (F 2, 271 = 13.05, P < 0.0001), mean (F 2, 271 = 6.019, P = 0.0028) and width (F 2, 271 = 1.695, P = 0.1856).

**Table 2:** The variables of the IUTR structure and the adjusted R^2^ values for the locomotor activity elicited by saline, cocaine and caffeine that includes vehicle pretreatment.

|  | saline | cocaine | caffeine |
| --- | --- | --- | --- |
| Adjusted R<sup>2</sup> | 0.8960 | 0.6669 | 0.7990 |
| Amplitude | 749.7223 ± 57.8881 | 1819.5871 ± 129.0375 | 1159.4926 ± 45.0063 |
| Mean | 47.5961 ± 1.4693 | 70.8736 ± 2.3861 | 73.9730 ± 1.7133 |
| width | 12.8420 ± 1.2797 | 25.8781 ± 3.2254 | 31.1834 ± 2.6770 |

When we compared the locomotor activity IUTR structures of cocaine and caffeine (excluding saline), we also detected significant differences (F 3, 190 = 5.544, P = 0.0011, Fig 4B). At the level of the individual variables, the results were as follows: amplitude (F 1, 190 = 12.49, P = 0.0005), mean (F 1, 190 = 0.2985, P = 0.5855) and width (F 1, 190 = 0.5177, P = 0.4727). In summary, under vehicle pretreatment, the QSCAn model detected differences in the IUTR structure of cocaine versus caffeine-induced locomotor activity (driven by differences in the amplitude variable).

### Cis-flupenthixol pretreatment: saline versus cocaine versus caffeine

For experiments that included cis-flupenthixol pretreatment (Fig 4C-D), Two-way repeated measures ANOVA with factors drug (saline, cocaine, caffeine) and time (14 levels) revealed the following: drug × time interaction (F 6.005, 51.04 = 2.761, P = 0.0211), no main effect of drug (F 2, 17 = 3.205, P = 0.0659), and a main effect of time (F 3.002, 51.04 = 11.25, P < 0.0001).

Tukey’s post hoc test revealed that the time course for saline and cocaine were different only at time = 60 min (* in Fig 4C). Tukey’s post hoc test revealed differences between saline versus caffeine at time = 60 min, 80 min and 85 min. There were no differences between cocaine and caffeine at all/any time points.

We wanted to compare the locomotor activity time course of cocaine and caffeine (without saline) under cis-flupenthixol pretreatment. Two-way repeated measures ANOVA revealed the following results: no psychostimulant × time interaction (F 2.868, 34.41 = 1.589, P = 0.2112), no main effect of psychostimulant (F 1, 12 = 0.06802, P = 0.7987), but a main effect of time (F 2.868, 34.41 = 9.751, P = 0.0001). In summary, under cis-flupenthixol pretreatment, the current model did not detect any differences in the time course of caffeine and cocaine-related locomotor activity.

For QSCAn model, we fitted the IUDR: a table of the variables of the IUTR structure and the adjusted R^2^ values are shown in Table 3. A comparison between the IUDR structures for saline, cocaine and caffeine (following dopamine receptor blockade) showed significant differences (F 6, 257 = 13.29, P < 0.0001, Fig 4D). For the variables comparisons, we obtained the following results: amplitude (F 2, 257 = 8.920, P = 0.0002), mean (F 2, 257 = 3.276, P = 0.0394) and width (F 2, 257 = 1.712, P = 0.1826).

**Table 3:** The variables of the IUTR structure and the adjusted R^2^ values for the locomotor activity elicited by saline, cocaine and caffeine that includes cis-flupenthixol pretreatment.

|  | saline | cocaine | caffeine |
| --- | --- | --- | --- |
| Adjusted R<sup>2</sup> | 0.9356 | 0.8343 | 0.7708 |
| Amplitude | 494.0097 ± 34.8402 | 1304.5616 ± 104.9010 | 840.1467 ± 51.2095 |
| Mean | 45.1533 ± 1.0408 | 58.3017 ± 1.6381 | 60.4199 ± 2.3372 |
| width | 10.9283 ± 0.8718 | 14.7882 ± 1.7140 | 24.6986 ± 2.5775 |

When we compared the structures of the locomotor activity (following cis-flupenthixol administration) of cocaine and caffeine (excluding saline), we detected that they had no differences in the overall structure (F 3, 176 = 1.853, P = 0.1393, Fig 4D). For variables comparisons, the results were as follows: amplitude (F 1, 176 = 5.270, P = 0.0229), mean (F 1, 176 = 0.1647, P = 0.6853) and width (F 1, 176 = 1.665, P = 0.1986). In summary, under cis-flupenthixol pretreatment, the QSCAn model detected differences in the structure (amplitude) of the time course of caffeine versus cocaine-related locomotor activity.

### Vehicle versus cis-flupenthixol: saline

For the saline group (Fig 5A-B), Two-way repeated measures ANOVA with factors pretreatment (vehicle, cis-flupenthixol) and time (14 levels) revealed the following: pretreatment × time interaction (F 2.581, 12.91 = 1.902, P = 0.1834), a main effect of pretreatment (F 1, 5 = 10.94, P = 0.0213), and a main effect of time (F 3.262, 16.31 = 16.63, P < 0.0001). However, Tukey’s post hoc tests revealed no significant differences between saline-induced locomotor activity for vehicle versus cis-flupenthixol at all/any time points (Fig 5A).

**Fig 5:**
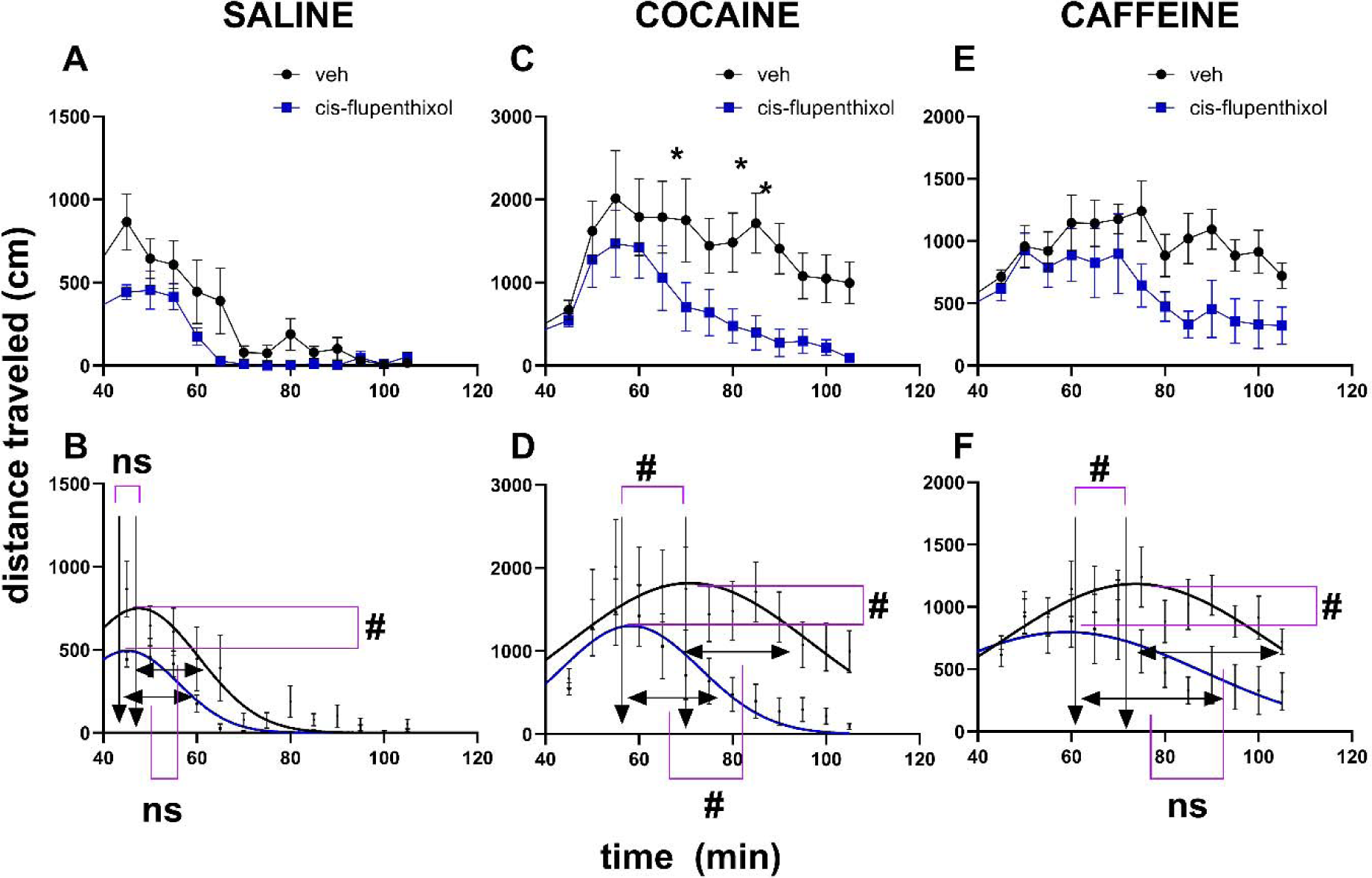
The Quantitative Structure of Curve Analytical (QSCAn) model was a more effective tool for the purposes of distinguishing the effects of cis-flupenthixol versus vehicle on locomotor activity due to saline, cocaine and caffeine. Black, red, and gray symbols and lines represent saline, cocaine, and caffeine treatment groups, respectively. A, C and E represent, respectively, current model analysis of locomotor activity to compare the effects of vehicle and cis-flupenthixol on saline, cocaine and caffeine. B, D and F represent, respectively, QSCAn model analysis of locomotor activity to compare the effects of vehicle and cis-flupenthixol on saline, cocaine and caffeine. For the saline group, Two-way repeated measures ANOVA revealed no significant differences between the effects of vehicle versus cis-flupenthixol on saline-induced locomotor activity at all/any time points (A). On the other hand QSCAn model analysis of the same showed significant differences between the IUTR structure amplitude (B). For the cocaine group Two-way repeated measures ANOVA revealed a pretreatment × time interaction (P = 0.0506) with significant differences between vehicle-cocaine and cis-flupenthixol-cocaine at time points 70, 85 and 90 min (* in C). On the other hand QSCAn model analysis of the effects of vehicle versus cis-flupenthixol revealed significant differences between the IUTR structure for all the variables (amplitude, mean, width, D). For the caffeine group, Two-way repeated measures ANOVA revealed no significant differences between vehicle versus cis-flupenthixol on caffeine-induced locomotor activity for at all/any time points (E). On the other hand, for the comparison between the IUTR structures for the same data, QSCAn model analysis detected differences in vehicle versus cis-flupenthixol for IUTR structure amplitude (P = 0.0028) and mean (P = 0.0011) but not width (P = 0.2751). In summary, QSCAn model was more effective than the current model in detecting differences in the effects of cis-flupenthixol relative to vehicle.

A comparison between the IUTR structure of the locomotor activity of vehicle-saline and cis-flupenthixol-saline showed significant differences (F 3, 162 = 9.697, P < 0.0001, Fig 5B). More detailed analysis of this IUTR structure comparison revealed the following results: amplitude (F 1, 162 = 13.97, P = 0.0003), mean (F 1, 162 = 1.009, P = 0.3166) and width (F 1, 162 = 0.8953, P = 0.3455).

### Vehicle versus cis-flupenthixol: cocaine

For the cocaine group (Fig 5C-D), Two-way repeated measures ANOVA revealed the following: pretreatment × time interaction (F 2.702, 18.92 = 3.210, P = 0.0506), a main effect of pretreatment (F 1, 7 = 10.23, P = 0.0151), and a main effect of time (F 1.526, 10.68 = 9.455, P = 0.0063). Tukey’s post hoc tests revealed significant differences between vehicle-cocaine and cis-flupenthixol-cocaine at time points 70, 85 and 90 min (* in Fig 5C).

A comparison between the IUTR structures for vehicle-cocaine and cis-flupenthixol-cocaine showed significant differences (F 3, 218 = 16.70, P < 0.0001, Fig 5D). More detailed analysis of this IUTR structure comparison revealed the following results: amplitude (F 1, 218 = 4.355, P = 0.0381), mean (F 1, 218 = 11.89, P = 0.0007) and width (F 1, 218 = 4.535, P = 0.0343).

### Vehicle versus cis-flupenthixol: caffeine

For the caffeine group (Fig 5E-F), Two-way repeated measures ANOVA revealed the following: no pretreatment × time interaction (F 3.802, 19.01 = 2.015, P = 0.1354), a main effect of pretreatment (F 1, 5 = 8.802, P = 0.0313), and a main effect of time (F 2.696, 13.48 = 6.015, P = 0.0092). However, Tukey’s post hoc tests revealed no significant differences between vehicle versus cis-flupenthixol on caffeine-induced locomotor activity for at all/any time points (Fig 5E).

A comparison between the IUTR structures for vehicle-caffeine and cis-flupenthixol-caffeine showed significant differences (F 3, 162 = 14.32, P < 0.0001, Fig 5F). More detailed analysis of this IUTR structure revealed the following results: amplitude (F 1, 162 = 9.193, P = 0.0028), mean (F 1, 162 = 11.04, P = 0.0011) and width (F 1, 162 = 1.199, P = 0.2751).

In summary, the QSCAn model was more effective than the current model in distinguishing cocaine versus caffeine-induced locomotor activity in the absence and presence of dopamine receptor blockade (Fig 4). Additionally, the QSCAn model was more effective than the current model in distinguishing vehicle versus cis-flupenthixol effects on cocaine- and caffeine-induced locomotor activity (Fig 5). A summary of our results are shown in Table 4.

**Table 4:** Summary of results from model comparison: current model versus QSCAn model.

|  | Current model | QSCAN model | Reference |
| --- | --- | --- | --- |
| Cocaine versus caffeine locomotor activity in absence of dopamine receptor blockade | No significant difference | Significant differences in IUTR structure, significant differences in amplitude | Fig 4A versus Fig 4B |
| Cocaine versus caffeine locomotor activity in presence of dopamine receptor blockade | No significant difference | Significant differences in IUTR structure, significant differences in amplitude | Fig 4C versus Fig 4D |
| Vehicle versus cis-flupenthixol (saline-induced locomotor activity) | No significant differences | Significant differences in IUTR structure, significant differences in amplitude | Fig 5A versus Fig 5B |
| Vehicle versus cis-flupenthixol (cocaine-induced locomotor activity) | Significant differences at 3 time points | Significant differences in IUTR structure, significant differences in amplitude, mean and width | Fig 5C versus Fig 5D |
| Vehicle versus cis-flupenthixol (caffeine-induced locomotor activity) | No significant differences | Significant differences in IUTR structure, significant differences in amplitude, mean but not width | Fig 5E versus Fig 5F |

## Discussion

We have developed a new QSCAn model as an advancement over the current model for the assessment of locomotor activity structure (Fig 2). Our hypothesis was that a new QSCAn model would be more effective than the current model in distinguishing differences between cocaine and caffeine dopamine-mediated locomotor activity. We designed experiments (Fig 3) to test our hypothesis. We determined that QSCAn model was more sensitive than the current model in distinguishing cocaine versus caffeine-induced locomotor activity in the absence and presence of dopamine receptor blockade (Fig 4-5). Our findings are summarized in Table 4. We confirmed our hypothesis.

There are many firsts in this study. We are the first to define and analyze locomotor activity as an IUTR curve structure. We are the first to identify an IUTR variable – width – that may reveal differences between cocaine versus caffeine. This is important because this variable that distinguishes cocaine versus caffeine locomotor activity is unrelated to the variables that are commonly used in the field. The new model does not mean that we should get rid of the old model rather we can combine both current and QSCAn models for more detailed analysis of locomotor activity going forward.

There are several implications for this study. By showing that we can differentiate between cocaine versus caffeine, this new model may be potentially useful in distinguishing the behavioral activity structures of different drugs including 1) different psychostimulants (cocaine versus amphetamines), 2) similar psychostimulants, 3) different classes of drugs (psychostimulants versus opioids) as long as these drugs can increase similar behavior over time. By showing that we can differentiate between the effects of vehicle and dopamine receptor antagonist, this new model may be potentially useful in distinguishing the pharmacological activity on behavioral structure of 1) different doses of the same drug, 2) different drugs of the same class, 3) different classes of drugs, and 4) different intervention strategies aimed at modulating behavior.

There are some limitations to this study that have to do with biological sex and drug doses. With respect to biological sex, we used only male rats, and it is possible that there will be biological sex differences in our data. However, we do not think this will be the case as we have shown in a study in which we employed male and female Sprague Dawley rats that biological sex differences are less significant than individual differences, at least with regards to psychostimulant-induced dopamine-mediated locomotor activity (Tigano and Job, 2025). Notwithstanding, it will be important to replicate our findings in female Sprague Dawley rats. With respect to drug dose: we used only one dose of cocaine (10 mg/kg i.p), one dose of caffeine (20 mg/kg i.p) and one dose of cis-flupenthixol (0.2 mg/kg i.p) and our results may not necessarily extend to other doses of these drugs. We have provided a rationale for every dose of drug that we used (see Methods section for dose consideration rationale). To address this limitation, we plan to conduct similar experiments in the future using different doses of cocaine, caffeine and cis-flupenthixol.

## Conclusion

Although caffeine and cocaine both elicit dopamine-dependent locomotor activity, conventional analytical approaches fail to resolve qualitative differences in the resulting behavioral output. Using the QSCAn model, we identified novel variables that distinguish between cocaine- and caffeine-induced locomotor activity under both baseline conditions and dopamine receptor blockade. These findings demonstrate that, despite shared involvement of dopamine signaling, caffeine and cocaine produce mechanistically distinct locomotor activity profiles. Collectively, this work establishes QSCAn as a sensitive analytical framework that advances behavioral phenotyping beyond existing models and enables differentiation of psychostimulant-induced behaviors that may appear similar using traditional measures.

## Acknowledgement

The authors wish to acknowledge Dr. Michael J Kuhar in whose lab all of the experiments were carried out. All authors (JM, KA and MOJ) contributed to data analysis and to the writing of the manuscript. MOJ designed and conducted behavioral experiments and statistical analysis.

## Disclosures

The authors declare no conflicts of interest with respect to research, authorship and publication of this article.

## Funding

The work described was conducted at the Yerkes National Primate Research Center of Emory University. This project was funded by the National Center for Research Resources [P51RR165] and supported by the Office of Research Infrastructure Programs/OD [P51OD11132]. It was also supported by the National Institutes of Health [DA15040, to MJK] and the Georgia Research Alliance. The development of the new analytical model was supported by Rowan University startup funds to MOJ. These funding sources had no role in the direction of this research or publication.

## Notes

### Competing Interest Statement

The authors have declared no competing interest.

### Summary of Updates

The Figure 5 in the previous version was not the right figure. This revision is to add the right Figure 5.

